# The microtubule GTP-tubulin cap size is modulated during cell division

**DOI:** 10.64898/2026.01.13.699367

**Authors:** Anna C. Cassidy, Dylan T. Burnette, Marija Zanic

## Abstract

Microtubule dynamics change during cell division to enable rapid microtubule network remodeling. The switching from microtubule growth to shrinkage is attributed to the loss of a stabilizing GTP-cap structure at the growing microtubule end. The size of the GTP-cap is a result of a balance between GTP-tubulin addition to the microtubule end and subsequent GTP-hydrolysis in the microtubule lattice. Whether the cell-cycle-dependent changes in microtubule dynamics are supported by concurrent modulation of the stabilizing GTP-cap size is not known. Here, we use high spatiotemporal resolution live-cell imaging of EB1, an established marker for the GTP-cap, to directly determine the relationship between GTP-cap size and microtubule growth rate throughout the cell cycle. Our data reveal that GTP-cap size for matching growth rates is reduced during mitosis. Comparison of EB1 comets on astral versus spindle microtubules reveals that the scaling between the GTP-cap size and microtubule growth rate is not spatially regulated in mitosis. We find that these regulatory patterns are conserved across epithelial cells from two different species. Taken together, our findings reveal modulation of GTP-cap size as a cell-cycle–regulated mechanism for tuning microtubule stability.

**Significance Statement:** Microtubule dynamics are altered during the cell cycle to enable rapid microtubule network remodeling and accurate chromosome segregation. By comparing EB1 comets on microtubule ends during different cell cycle stages, the authors find that microtubule GTP-cap size is subject to global differential regulation during specific cell cycle stages. These results identify modulation of microtubule stabilizing GTP-cap size as a previously underappreciated, cell-cycle–regulated mechanism for tuning microtubule stability throughout the cell cycle.

## INTRODUCTION

Microtubules are dynamic cytoskeletal polymers essential for myriad cellular processes including cell division, cell motility, and intracellular transport (Alberts *et al*., 2002). During cell division, the mitotic spindle–composed of microtubules–must accurately divide the sister chromatids into two new daughter cells. To form the mitotic spindle, the interphase microtubule cytoskeleton is completely disassembled as the cell enters mitosis (Inoué, 1964; Inoué and Sato, 1967). The swift remodeling of the microtubule cytoskeletal network architecture to form the mitotic spindle, along with a diverse range of other cellular functions, is made possible by the modulation of microtubule dynamics. Indeed, disruption of microtubule dynamics during mitosis blocks cell cycle progression; this finding serves as the basis for major cancer therapeutics which function by directly altering microtubule dynamics via stabilization or destabilization of microtubule polymers (Jordan and Wilson, 2004).

A defining characteristic of microtubule dynamics is dynamic instability–the intrinsic ability of microtubules to rapidly switch between phases of growth and shrinkage (Mitchison and Kirschner, 1984; Kirschner and Mitchison, 1986). Microtubule dynamic instability is driven by the GTPase activity of tubulin. Specifically, microtubules grow via the addition of GTP-bound αβ-tubulin heterodimers to the growing microtubule end (Weisenberg *et al*., 1968; Weisenberg, 1972). Once incorporated into the lattice, the β-tubulin subunit hydrolyzes GTP (Jacobs *et al*., 1974; Kobayashi, 1975; Weisenberg *et al*., 1976; David-Pfeuty *et al*., 1977; Carlier and Pantaloni, 1982) introducing a destabilizing conformational change, and yielding an inherently unstable microtubule lattice (Carlier *et al*., 1984). However, GTP hydrolysis does not occur immediately following heterodimer addition (Carlier and Pantaloni, 1981), and is triggered only after the heterodimer is capped by the next incoming subunit (Carlier *et al*., 1987; Nogales *et al*., 1998). This delay in GTP hydrolysis results in a ‘GTP-cap’, a stabilizing region at the growing microtubule end (Mitchison and Kirschner, 1984). Loss of the stabilizing GTP-cap triggers the onset of catastrophe – the transition between microtubule growth and shrinkage (Kirschner and Mitchison, 1986; Drechsel and Kirschner, 1994; Caplow and Shanks, 1996). Thus, the properties of the GTP-cap have direct implications on the regulation of microtubule dynamics.

Efforts to investigate the spatiotemporal features of the GTP-cap were greatly facilitated by the discovery that EBs, a family of autonomous microtubule tip-tracking proteins, are sensitive to the nucleotide state of tubulin in the microtubule polymer (Zanic *et al*., 2009; Maurer *et al*., 2012; Zhang *et al*., 2015; Duellberg *et al*., 2016; Roostalu *et al*., 2020). EBs preferentially bind the stabilizing GTP-cap, resulting in a comet-like localization at growing microtubule ends, reflecting the exponential decay of the GTP-bound tubulin state along the microtubule lattice due to the GTP hydrolysis. GTP-cap size is modulated by two key parameters: rate of addition of GTP-tubulin dimers to the growing microtubule end and the rate of GTP-hydrolysis within the polymer. Using EB localization as a proxy for the GTP-cap, prior investigations have revealed that EB comet size scales linearly with microtubule growth rate both *in vitro* and in cells (Bieling *et al*., 2007, 2008; Seetapun *et al*., 2012; Duellberg *et al*., 2016; Rickman *et al*., 2017; Vemu *et al*., 2017; Roth *et al*., 2018; Strothman *et al*., 2019; Farmer*, Arpağ* *et al*., 2021; Urazbaev *et al*., 2021; Liao *et al*., 2022; Cassidy*, Farmer* *et al*., 2024), consistent with a fixed GTP-hydrolysis rate. Moreover, a direct comparison of EB comets *in vitro* to those in cells revealed that the relationship between GTP-cap size and microtubule growth rate is the same in interphase cells as *in vitro*, suggesting that the tubulin GTP-hydrolysis rate is not differentially regulated in that cellular context (Cassidy*, Farmer* *et al*., 2024). However, previous studies have not investigated the relationship between the GTP-cap size and the microtubule growth rate throughout the cell cycle.

Microtubule dynamics and network architecture change significantly during cell division (Verde *et al*., 1992; Rusan *et al*., 2001; Yamashita *et al*., 2015), supported by changes in the expression levels and post-translation modifications of numerous microtubule regulatory proteins (Ferreira *et al*., 2014, 2018; Vicente and Wordeman, 2019). Thus, we hypothesized that GTP-cap size may be differentially modulated during mitosis as an additional mode of regulation to achieve altered microtubule dynamics required for successful cell division. Here, we use high-spatiotemporal-resolution imaging of EB1 comets in two different mammalian epithelial cell types and across six distinct points in the cell cycle to investigate if GTP-cap size is differentially modulated during cell division.

## RESULTS

### Microtubules display overlapping growth rates throughout the cell cycle

To directly compare the relationship between the GTP-cap size and microtubule growth rate at different points in the cell cycle, we imaged porcine kidney epithelial (LLC-PK1) cells stably expressing EB1-GFP and transiently expressing mCherry-H2B, a fluorescently-tagged histone protein. We used LLC-PK1 cells (Piehl and Cassimeris, 2003), an established model for imaging microtubule dynamics throughout the cell cycle, since they remain flat throughout mitosis (Gorbsky *et al*., 1988; Rusan *et al*., 2001; Wadsworth *et al*., 2005). Furthermore, we verified that variations in EB1-GFP expression levels among the stably expressing LLC-PK1 cells do not influence microtubule dynamics, consistent with previous reports (Piehl and Cassimeris, 2003; Cassidy*, Farmer* *et al*., 2024) (Figure S1). Transient transfection with mCherry-H2B allowed us to visualize cellular DNA organization and determine different cell cycle stages (Figures 1A-F top images, Video 1) (Kanda *et al*., 1998). We selected six distinct points in the cell cycle for this study: interphase, prophase, prometaphase, metaphase, anaphase, and two daughter cells following cytokinesis.

**Figure 1.**
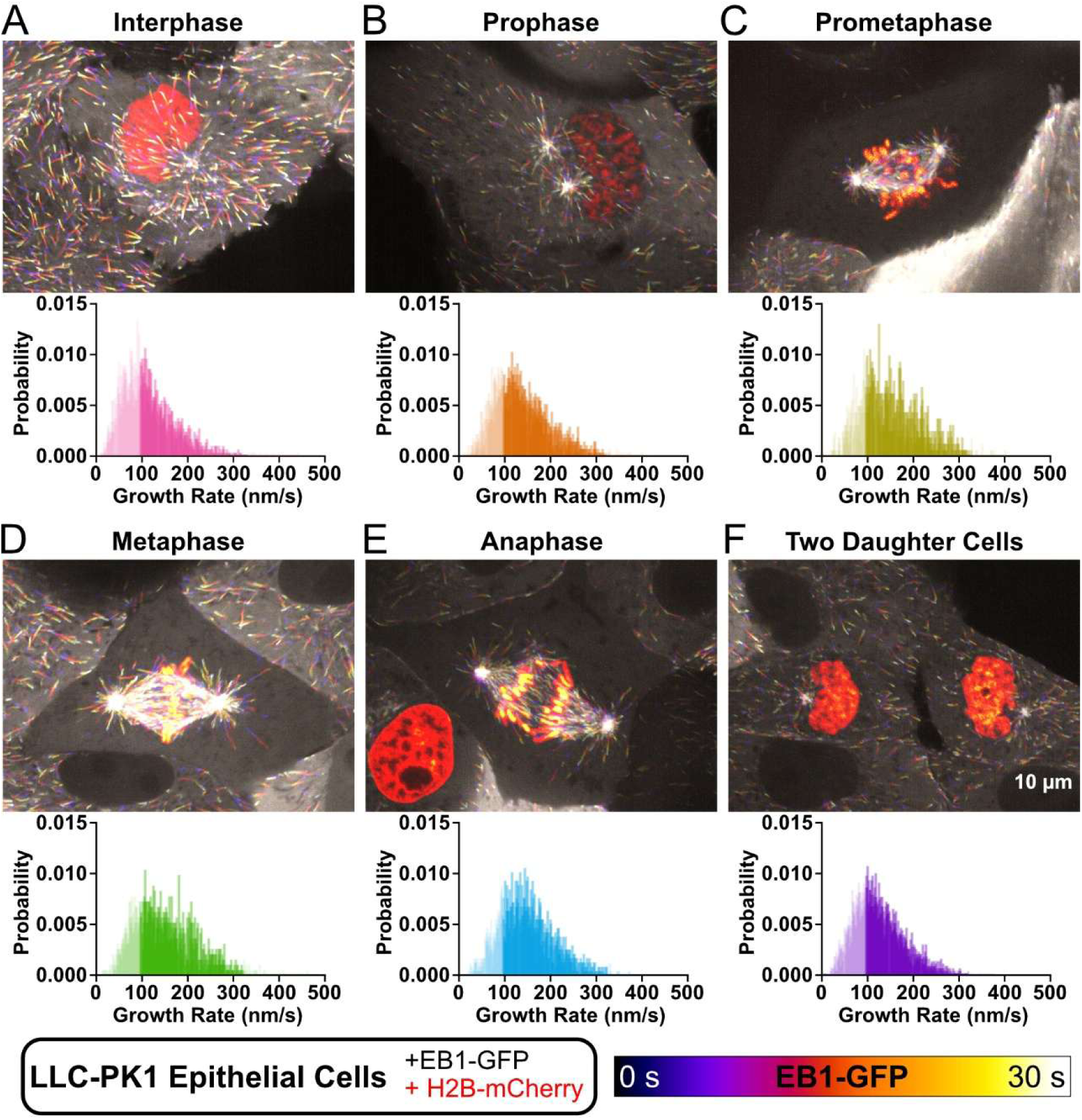
Microtubules display overlapping growth rates throughout the cell cycle. (A-F) Top, representative images of temporally-colored maximum intensity projection of EB1-GFP comets over time in LLC-PK1 cells stably expressing EB1-GFP and transiently expressing mCherry-H2B during a 30-second movie. Chromosomes are visualized at the starting timepoint via acquisition of a single frame of mCherry-H2B signal shown in orange. Cells were classified into different cell cycle stages based on chromosome organization: (A) interphase, (B) prophase, (C) prometaphase, (D) metaphase, (E) anaphase, and (F) two daughter cells. Bottom, probability distribution of microtubule growth rates for each cell cycle stage. Growth rates are binned into 1 nm/s bins. Dark shaded bars denote microtubules within a growth rate range used for subsequent analysis. Data were acquired from 3 separate cells per day over 3 days for each stage (N=9 cells per stage). Probability is displayed as a ratio (out of 1).

Given that microtubule dynamics change throughout the cell cycle (Rusan *et al*., 2001; Yamashita *et al*., 2015), we first wanted to characterize the range and distribution of microtubule growth rates for all cells in our dataset to ensure that there was sufficient overlap between each of the selected cell cycle timepoints. This was essential, as we were interested in directly comparing EB1 comet lengths for matched microtubule growth rates in order to eliminate any potential differences in growth rate as a confounding variable in our investigations (Farmer*, Arpağ* *et al*., 2021; Cassidy*, Farmer* *et al*., 2024). Our analysis confirmed that microtubule growth rates change during different stages of the cell cycle, as expected (Piehl and Cassimeris, 2003; Yamashita *et al*., 2015). Nevertheless, a substantial overlap in growth rates was observed across all stages (Figure 1). Therefore, it is possible to directly compare EB1 comets between different cell cycle stages across a range of matched microtubule growth rates (Figure 1A-F bottom graphs).

### GTP-cap size is reduced during mitosis

To determine whether the GTP-cap size is modulated from one cell cycle stage to another, we measured the lengths of EB1-GFP comets as a function of microtubule growth rate for six selected cell cycle stages. We found that all cell cycle stages had microtubules growing at rates up to 320 nm/s, which set the upper limit of our range of matched growth rates. Furthermore, microtubules growing at rates above 100 nm/s displayed resolvable EB1-GFP comets in all investigated cell cycle stages, setting the lower limit of our growth-rate range. To quantify the size of the GTP-cap on each microtubule, we determined the decay length of the exponential decay function that was fit to the EB1-GFP comet intensity profile (see *Materials and Methods*) (Farmer*, Arpağ* *et al*., 2021; Cassidy*, Farmer* *et al*., 2024). In this way, our analysis reflected the spatial decay of the GTP-cap along the microtubule, ensuring that our measurements were independent of the absolute intensity of EB1-GFP localization on any given microtubule.

As previously reported, EB1 comet lengths scaled linearly with microtubule growth rate in interphase cells (Figure 2A) (Urazbaev *et al*., 2021; Liao *et al*., 2022; Cassidy*, Farmer* *et al*., 2024). We also observed linear scaling between microtubule growth rate and EB1 comet length across all other examined cell cycle stages (Figure 2B–F). However, when these data were fit to a linear regression, we found that this relationship—specifically the slope—is not conserved across stages (Figure 2G; p-value < 0.0001). Instead, for matched microtubule growth rates, EB1 comet length gradually decreased from interphase through prophase to prometaphase. EB1 comets in metaphase displayed intermediate lengths, larger than prometaphase, yet smaller than interphase and prophase. Anaphase comets were reduced in size relative to metaphase, displaying lengths similar to prometaphase comets. Finally, investigation of EB1 comets in post-cytokinetic cells showed that EB1 comet lengths increased upon mitotic exit, suggesting a return to the interphase preset (Figure 2H).

**Figure 2.**
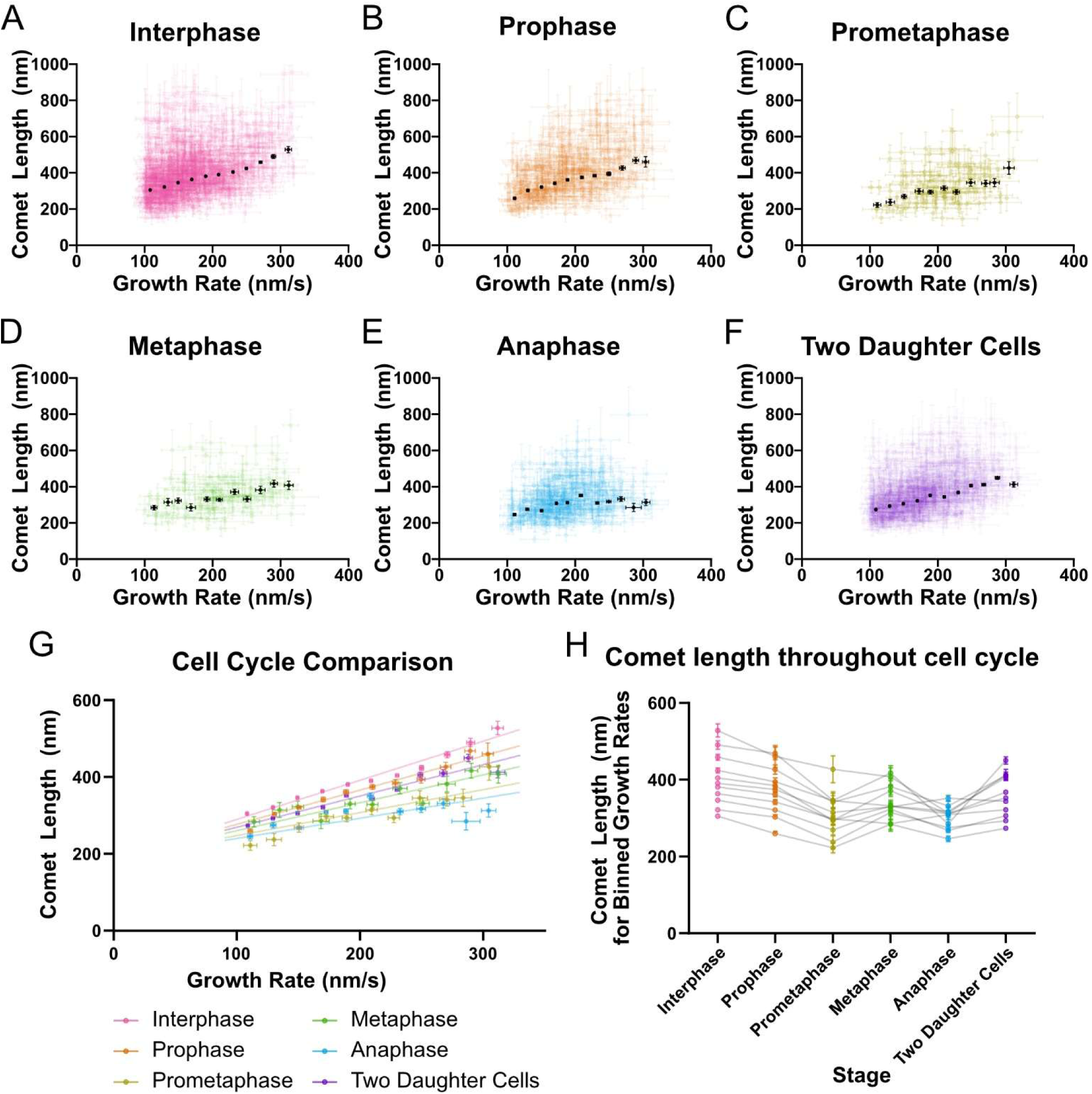
The scaling between the EB1 comet length and the microtubule growth rate is modulated during the cell cycle. (A-F) EB1 comet length as a function of microtubule growth rate for the same cells analyzed in Figure 1, bottom. Data were acquired from 3 separate cells per day over 3 days for each stage (N=9 cells per stage). Colored points are the mean growth rate ± 95% CI and EB1 comet length for individual growth events lasting ≥ 4 s. Black dots represent weighted averages with weighted errors of 95% CI for each 20 nm/s bin. (A) N = 1,231 growth events. (B) N = 471 growth events. (C) N = 120 growth events. (D) N = 133 growth events. (E) N = 402 growth events. (F) N = 1,013 growth events. (G) Binned weighted average data points for each cell-cycle stage (black points in A-F) were fit using unweighted least-squares linear regression. (H) Average EB1 comet length for matched growth rate bins (data from G) connected across stages by grey lines.

To assess whether subsets of cell cycle stages could be described by a common linear model, we repeated the linear regression analysis using grouped stages. Only prometaphase and anaphase could be fit by a single linear model (p-value = 0.19). Notably, the relationship between EB1 comet length and microtubule growth rate observed in interphase was statistically significantly different from that of all other stages, demonstrating that the GTP-cap size is differentially modulated as cells enter and progress through the cell cycle.

### GTP-cap size is not spatially regulated in metaphase or anaphase

When we investigated the relationship between the GTP-cap size and microtubule growth rate in different cell cycle stages, we were agnostic about the spatial location of the analyzed EB1 comets in each cell. After finding cell-cycle-dependent differences in the relationship between these parameters, we wondered if the GTP-cap size was also subject to differential spatial regulation within a single cell. Subcellular spatial regulation of microtubule dynamics has been demonstrated in many different contexts, including mitotic (Stumpff *et al*., 2012; Fukuda *et al*., 2014; Yamashita *et al*., 2015; Walczak *et al*., 2016), migratory (Etienne-Manneville *et al*., 2005; Wittmann and Waterman-Storer, 2005; Trogden and Rogers, 2015), and neuronal cells (Tan *et al*., 2019). In mitotic cells, there is robust evidence for spatial regulation of astral versus spindle microtubule dynamics spanning from modulation of microtubule stability (Stumpff *et al*., 2012; Fukuda *et al*., 2014; Walczak *et al*., 2016) to differences in microtubule growth rates (Yamashita *et al*., 2015).

For our study, we decided to specifically look at spindle versus astral microtubule GTP-cap sizes in metaphase and anaphase cells where these microtubule populations could be visually distinguished from each other (Figure 3A) (Walczak *et al*., 2016). To accomplish this, we revisited our metaphase and anaphase datasets and classified each analyzed comet as either astral (Figure 3B,E) or spindle (Figure 3C,F). When comparing the spatial distribution of microtubule growth rates in each stage, we found that metaphase astral microtubules (Figure 3B) generally had faster microtubule growth rates compared to spindle microtubules (Figure 3C), in agreement with previous reports (Yamashita *et al*., 2015). Surprisingly, fitting these data to a linear regression revealed that the relationship between the EB1comet length and the microtubule growth rate is not spatially regulated in metaphase (Figure 3D, slope of 0.68 ± 0.11 s [95% confidence interval, CI] for spindle vs. 0.78 ± 0.19 s [95% CI] for astral, p-value = 0.27). Similarly, while anaphase comets for a given growth rate were overall smaller than those observed in metaphase, they also did not display any spatial variation in the relationship between the growth rate and the comet length (Figure 3G, slope of 0.56 ± 0.11 s [95% CI] for spindle vs. 0.51 ± 0.10 s [95% CI] for astral, p-value = 0.43). This finding suggests that the reduction in GTP-cap size at matched microtubule growth rates during mitosis is regulated globally rather than locally.

**Figure 3.**
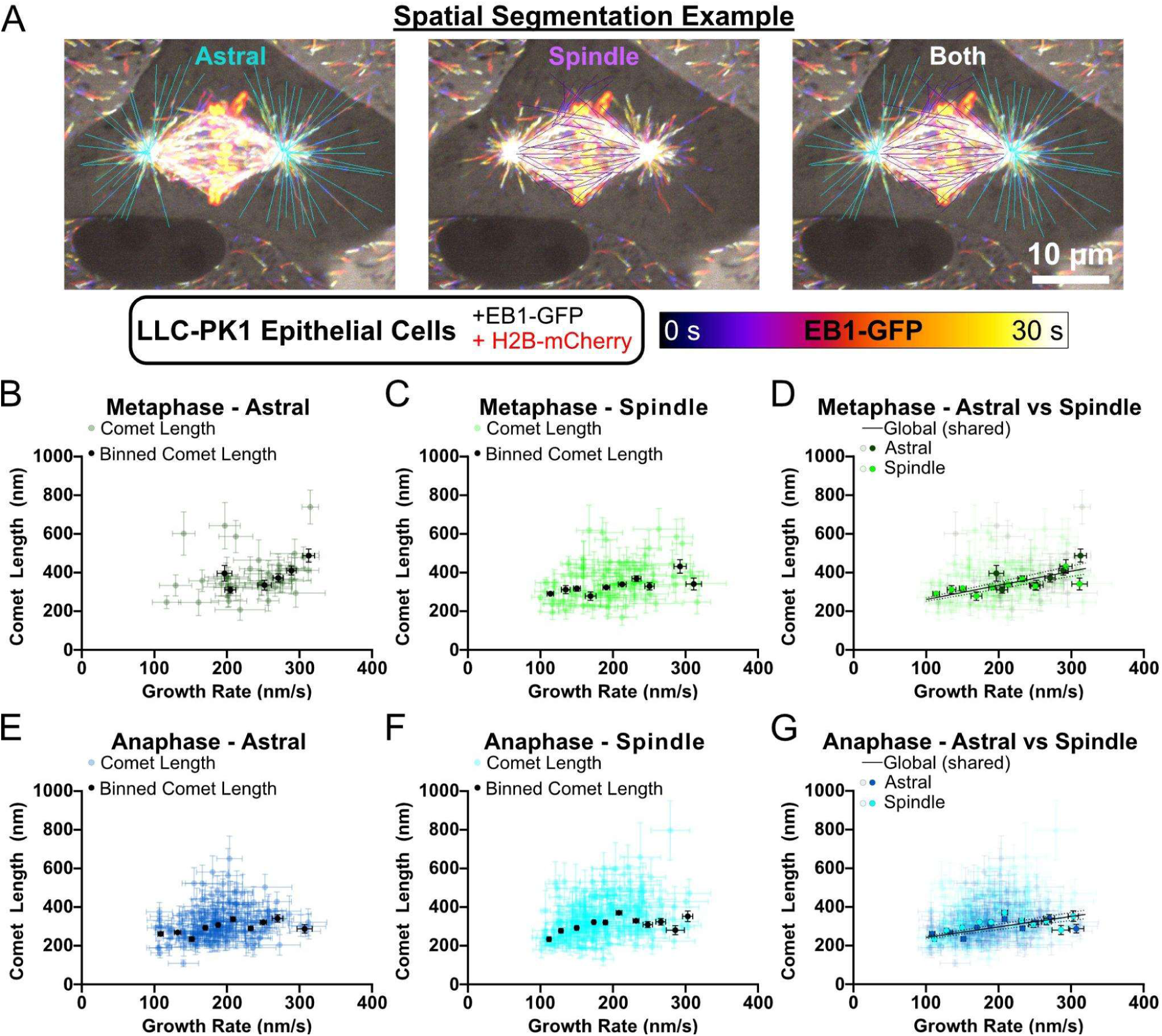
EB1 comet lengths of astral and spindle microtubules are not differentially regulated in mitosis. (A) Example illustrating the spatial designation of EB1 comets as belonging to astral or spindle microtubules. (B–C, E–F) Spatial segmentation of EB1 comet length as a function of microtubule growth rate during metaphase and anaphase for the same cells analyzed in Figure 2. EB1 comets localized to the tips of astral (B, E) or spindle (C, F) microtubules are shown. Colored points represent the mean growth rate and EB1 comet length for individual growth events lasting ≥ 4 s, with ±95% confidence intervals (CI). Individual events were binned in 20 nm/s intervals. Black points represent weighted averages with weighted errors of 95% CI for each bin containing ≥ 3 growth events. (B) N = 42 growth events. (C) N = 91 growth events. (E) N = 188 growth events. (F) N = 214 growth events. (D, G) Binned weighted averaged data points for astral and spindle microtubules in each cell cycle stage (black points in B–C and E–F) were fit using unweighted least-squares linear regression. Dotted lines denote the 95% CI of the fits. (D) p-value = 0.27. (G) p-value = 0.43.

### Changes in GTP-cap size are conserved across cell types

Based on our data supporting global regulation of the GTP-cap size during the cell cycle, we sought to determine whether this mechanism is conserved across species. To that end, we selected PtK1 cells, a potoroo kidney epithelial cell line, stably expressing EB1-GFP (Tirnauer *et al*., 2002) because, like LLC-PK1 cells, PtK1 cells remain flat throughout mitosis (Wadsworth *et al*., 2005) and are an established model for imaging microtubule dynamics during mitosis (Stout *et al*., 2006; Dumont and Mitchison, 2009). As before, we assigned cell cycle stage based on H2B localization (Kanda *et al*., 1998), imaged EB1 comets in PtK1 cells over time, and quantified EB1 comet length and microtubule growth rate using the same analysis pipeline (see *Materials and Methods*). We found that EB1 comet length scaled linearly with microtubule growth rate in interphase PtK1 cells (Figure 4A), as well as across all other examined cell cycle stages (Figure 4B–F), consistent with our observations in LLC-PK1 cells. Furthermore, in agreement with LLC-PK1 data, the relationship between microtubule growth rate and GTP-cap size for different cell cycle stages could not be described by a single linear regression model (Figure 4G; p-value < 0.0001). For microtubules growing at matched growth rates, EB1 comets were generally smaller during mitosis, followed by an apparent “reset” upon mitotic exit (Figure 4H). In both investigated cell types, EB1 comet lengths were smallest relative to their corresponding matched growth rate during prometaphase and anaphase (Figure 4H).

**Figure 4.**
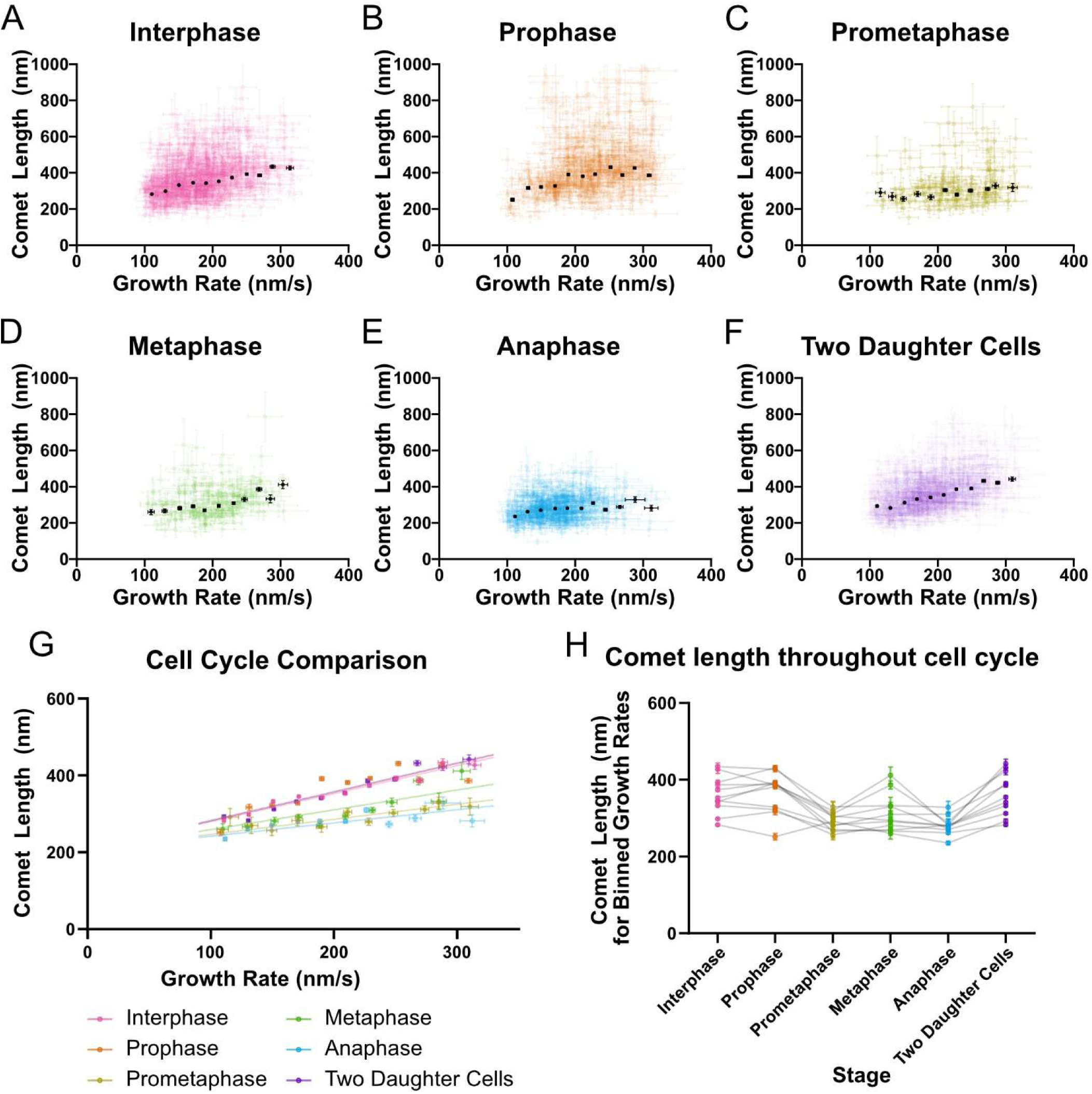
Changes in EB1 comet length *are* conserved across cell types. (A-F) EB1 comet length as a function of microtubule growth rate in PtK1 cells. Data were acquired from 3 separate cells per day over 3 days for each stage (N=9 cells per stage). Colored points are the mean growth rate ± 95% CI and EB1 comet length for individual growth events lasting ≥ 4 s. Black dots represent weighted averages with weighted errors of 95% CI for each bin for bins containing ≥ 3 growth events. (A) N = 1041 growth events. (B) N = 548 growth events. (C) N = 144 growth events. (D) N = 179 growth events. (E) N = 585 growth events. (F) N = 891 growth events. (G) Binned weighted average data points for each cell-cycle stage (black points in A-F) were fit using unweighted least-squares linear regression. (H) Average EB1 comet length for matched growth rate bins (data from G) connected across stages by grey lines.

Consistent with our results in LLC-PK1 cells, EB1 comets during prometaphase and anaphase in PtK1 cells could be described by a single linear regression (p-value = 0.17). Additionally, EB1 comets in PtK1 cells during interphase, prophase, and in the two daughter cells could also be described by a single linear model (p-value = 0.81). Metaphase remained statistically distinct from interphase, similar to our observations in LLC-PK1 cells. Taken together, these findings suggest that modulation of the GTP-cap size represents a general mechanism for mitotic-specific regulation of microtubule stability and dynamics, supporting extensive microtubule network remodeling that accompanies cell division.

## CONCLUSIONS

It has been long recognized that microtubule dynamics change throughout the cell cycle to support microtubule network remodeling in cell division (Inoué, 1964; Inoué and Sato, 1967). Whether these changes in dynamics are accomplished through modulation of the size of the stabilizing GTP-cap at the growing microtubule ends was not known. In this work, we systematically investigated the relationship between the GTP-cap size and microtubule growth rate throughout the cell cycle. We took advantage of the finding that EB proteins specifically recognize the nucleotide state of tubulin at growing microtubule ends, building on the extensive body of previous work that established EBs as the bona fide marker of the stabilizing GTP-cap (Zanic *et al*., 2009; Maurer *et al*., 2012; Zhang *et al*., 2015; Duellberg *et al*., 2016; Roostalu *et al*., 2020). We confirmed that the expression levels of the EB1-GFP reporter in our experiments did not affect microtubule dynamics (Fig S1). Furthermore, our method of measuring EB1-comet decay lengths, rather than absolute intensities of bound EB1-GFP, ensured that our measurements of the microtubule GTP-cap size were not affected by any potential cell-to-cell variability in the EB1-GFP fluorescence intensity. By directly comparing decay lengths of EB1 comets from microtubules with matched growth rates across six distinct cell cycle stages we found that the relationship between GTP-cap size and microtubule growth rate is globally altered in a cell-cycle-stage-dependent manner. Therefore, our results identify modulation of the microtubule GTP-cap size as a potentially powerful mechanism to tune microtubule stability throughout the cell cycle, supporting the dramatic reorganization of the microtubule cytoskeleton during mitosis and the need for both rapid turnover and precise control of microtubule attachments.

From a functional perspective, a reduction in the GTP-cap size at a given growth rate would be expected to increase catastrophe probability and enhance microtubule responsiveness to destabilizing cues (Duellberg *et al*., 2016; Roostalu *et al*., 2020; Farmer*, Arpağ* *et al*., 2021). Such a regime is particularly well suited to prometaphase and anaphase, which represent periods of heightened microtubule turnover and responsiveness. In prometaphase, rapid microtubule turnover facilitates error correction (Cimini *et al*., 2006; Bakhoum *et al*., 2009), while in anaphase, coordinated microtubule disassembly contributes to spindle elongation and chromosome segregation (Inoué and Salmon, 1995). In both stages, simply slowing growth would be insufficient or counterproductive; instead, reducing cap size at a given growth rate provides a mechanism to maintain fast dynamics while increasing turnover. Indeed, our results demonstrate that, for microtubules growing at matched growth rates, GTP-cap size is most significantly reduced in prometaphase and anaphase. By contrast, metaphase requires a temporary balance between stability and responsiveness to reinforce correct microtubule-kinetochore attachments (Zhai *et al*., 1995); this is consistent with the lesser degree of GTP-cap size reduction we observed in metaphase. Furthermore, this intermediate effect observed in metaphase suggests that GTP-cap size regulation is tunable, rather than simply switched on for all of mitosis. Taken together, our observation that the relationship between GTP-cap size and microtubule growth rate is altered within specific cell cycle stages, while growth rates themselves remain within overlapping ranges, suggests that mitosis introduces an additional regulatory layer: modifying the GTP-cap size without necessarily changing how fast microtubules grow.

Spatial regulation of microtubule dynamics is a well-established feature of many cellular contexts, including mitosis (Stumpff *et al*., 2012; Fukuda *et al*., 2014; Yamashita *et al*., 2015; Walczak *et al*., 2016). Interestingly, our investigation of the GTP-cap size showed no differences in astral versus spindle microtubules within a given mitotic stage. This result argues against spatially restricted mechanisms, such as localized force generation or compartmentalized MAP activity, as primary drivers of the GTP-cap size modulation. Instead, it suggests that mitotic regulation of the GTP-cap size reflects a global, cell-wide change that uniformly affects growing microtubule plus ends, independent of their spatial location or functional role within the spindle.

The size of the GTP-cap is set through a competition between the rates of soluble GTP-tubulin addition to growing microtubule ends and subsequent GTP-hydrolysis in the microtubule polymer. Thus, a faster tubulin incorporation is expected to increase the GTP-cap size, consistent with our findings that faster microtubule growth rates correlate with an increase in the GTP-cap size across all cell cycle stages and widely supported by the literature in vitro and in cells (Bieling *et al*., 2007, 2008; Duellberg *et al*., 2016; Rickman *et al*., 2017; Vemu *et al*., 2017; Roth *et al*., 2018; Strothman *et al*., 2019; Farmer*, Arpağ* *et al*., 2021; Urazbaev *et al*., 2021; Liao *et al*., 2022; Cassidy*, Farmer* *et al*., 2024). Conversely, an increase in the GTP-hydrolysis rate within the polymer is expected to shorten the GTP-cap, even if the microtubule growth rate is unchanged. Thus, the cell-cycle dependent modulation of the GTP-cap size for a given growth rate observed in our experiments could be explained by mitosis-specific regulation of the GTP-hydrolysis rate. Indeed, mitosis is characterized by widespread changes in the expression levels, localization, and post-translational modifications of microtubule-associated proteins which regulate microtubule dynamics (Ferreira *et al*., 2014, 2018; Vicente and Wordeman, 2019). To that end, mitotic MAP activity may accelerate GTP-tubulin turnover by promoting post-hydrolysis conformations, facilitating phosphate release, or otherwise shortening the EB-binding region.

Hydrolysis of GTP induces structural changes within the microtubule lattice, which may involve both compaction and twist in the arrangement of the tubulin dimers (Hyman *et al*., 1995; Nogales *et al*., 1998; Alushin *et al*., 2014; Zhang *et al*., 2018; LaFrance *et al*., 2022). Recent studies implicated a number of MAPs and motors in both inducing and recognizing either compacted or expanded conformations of the microtubule lattice (Zhang *et al*., 2017; Peet *et al*., 2018; Liu and Shima, 2023; Verhey and Ohi, 2023). To what extent these lattice conformations correlate with specific states in the tubulin GTPase cycle is not yet understood. In addition to directly controlling the GTP hydrolysis rate, MAPs can disrupt the structural integrity of the growing microtubule end, and thus the stabilizing GTP-cap. Indeed, several MAPs have been reported to induce splayed, curled or lagging protofilaments at the microtubule end (Vitre *et al*., 2008; Farmer*, Arpağ* *et al*., 2021; Gudimchuk and McIntosh, 2021; Atherton *et al*., 2022). Coupled with the increased mechanical loads on microtubule tips during mitosis, all of these activities could effectively result in a reduction of the stabilizing GTP-cap region recognized by EBs. Deciphering of the precise interplay of the multitude of cellular factors thus remains an ongoing challenge for future research. Nevertheless, our findings identify modulation of microtubule stabilizing GTP-cap size as a previously underappreciated, cell-cycle–regulated mechanism for tuning microtubule dynamics.

## ACKNOWLEDGEMENTS

We thank R. Ohi (University of Michigan) and S. Dumont (University of California, San Francisco), for kindly gifting us the LLC-PK1 EB1-GFP cells and the PtK1 EB1-GFP cells, respectively. A portion of the SDC microscopy imaging was performed using the Vanderbilt Cell Imaging Shared Resource (supported by NIH grants CA68485, DK58404, and EY08126). Use of the core equipment was supported in part by Vanderbilt Ingram Cancer Center Resource Share Scholarship (2023-4031515). We thank members of the Burnette laboratory (Vanderbilt University) for help with SDC. We thank the members of the Zanic laboratory for discussions and feedback. This work was supported by the National Institutes of Health grants R35GM119552 to MZ and R35GM125028 to DTB. The authors declare no competing financial interests.

## MATERIALS AND METHODS

### Cell culture

LLC-PK1 cells stably expressing EB1-GFP (a pig kidney epithelial cell line) were maintained in a 1:1 mixture of OptiMEM (REF 31985-070) and Ham’s F-10 medium (REF 11550), supplemented with 10% fetal bovine serum (FBS) and 1% Penicillin-Streptomycin. The cell line, originally developed by Piehl and Cassimeris (Piehl and Cassimeris, 2003), was kindly provided by Dr. Ryoma Ohi (University of Michigan). PtK1 cells stably expressing EB1-GFP (a long-nosed potoroo kidney epithelial cell line) were maintained in Ham’s F-12 medium (REF 11765) supplemented with 10% fetal bovine serum (FBS) and 1% Penicillin-Streptomycin. The cell line, originally developed by the Salmon and Mitchinson labs (Tirnauer *et al*., 2002), was kindly provided by Dr. Sophie Dumont (University of California, San Francisco). Cultures for both cell lines were kept at 37°C in a humidified atmosphere with 5% CO₂. Prior to imaging, the growth medium was exchanged for FluoroBrite DMEM supplemented with 10% FBS to optimize conditions for live-cell fluorescence microscopy.

### Transfection

Both LLC-PK1 and PtK1 cells were transiently transfected with H2B-mCherry-18, which was a gift from Dylan Burnette (originally, a gift from Michael Davidson (Addgene plasmid # 55055 ; http://n2t.net/addgene:55055 ; RRID:Addgene_55055)), using Lipofectamine3000 at a 1:2 DNA to Lipofectamine ratio. Cells were incubated with the plasmid and Lipofectamine3000 mix for ∼4 h at 37°C in a humidified atmosphere with 5% CO₂. After the 4-h incubation, the media was aspirated, and fresh media was added to the dishes.

LLC-PK1 cells were transiently transfected with 1 ug of H2B-mCherry-18. Approximately 24 hours prior to transient transfection, LLC-PK1 EB1-GFP cells were split and seeded at ∼15-20% confluency into two 32 mm #1.5 poly-L-lysine precoated glass bottom dishes. Cells were imaged ∼20–24 h after transient transfection.

PtK1 cells were transiently transfected with 4 ug of H2B-mCherry-18. 3-4 days prior to transient transfection, PtK1 cells were split and seeded at ∼40-60% confluency into a T25 flask. Following transfection, the cells were incubated for ∼24 hours. At this point, cells were split and seeded into a 32 mm #1.5 poly-L-lysine precoated glass bottom dish at 60% confluency. Cells were imaged ∼72 hours after transient transfection (∼48 hours after dishes were seeded).

### Determination of the effect of EB1 overexpression on microtubule dynamics

To quantify EB1-GFP expression levels, corrected total cell fluorescence (CTCF) was measured using FIJI/ImageJ (Schindelin *et al*., 2015; Rueden *et al*., 2017). A freehand region of interest (ROI) was drawn around each cell, along with a background ROI drawn in a region without detectable fluorescence. For each frame, the ROI area, mean gray value (average fluorescence intensity per pixel), and integrated density (total fluorescence, accounting for ROI size) were recorded. CTCF was then calculated using the formula:

*CTCF = Integrated Density (cell ROI) − (Area of cell ROI × Mean Gray Value of background ROI)*.

CTCF values across all frames were averaged to yield a single representative value per cell.

To robustly and agnostically measure microtubule growth rates at the single cell level we used *plusTipTracker* (Applegate *et al*., 2011), a MATLAB (vR2023b; MathWorks)-based image analysis package for detecting and tracking EB comets. *plusTipTracker* performs optimally on time-lapse image sequences acquired at 0.5–2 seconds per frame. Because our raw data were collected at a higher temporal resolution (0.14–0.16 s/frame), we generated temporally downsampled substacks to bring the frame interval within the optimal range. Specifically, every 7th frame was selected starting from frame 1, resulting in an effective frame rate of approximately 1–1.16 s/frame. After generating the substacks, the inverted cell ROI (from the CTCF analysis) was used as a mask to eliminate background and restrict comet detection to the cell of interest. Comets were detected using anisotropic Gaussian filtering with an alpha value of 0.01 and a minimum distance of 5 pixels to avoid duplicate identification. Tracking was performed using the “Microtubule Plus-End Dynamics” application with segment merging and splitting disabled to reduce tracking errors. The resulting tracks were analyzed using the “Microtubule Dynamics Classification” module.

### Imaging

Imaging was performed using two different Nikon SDC microscopes. Both SDC microscopes were equipped with a CSU-X1 (confocal scanner unit). For all imaging, a Tokai Hit objective heater was used to maintain the sample at 37°C and with 5% CO2. Images were acquired using NIS-Elements (Nikon) software. SDC enabled fast time-lapse imaging (∼7 fps) and high spatial resolution with the least photo damage–something we were particularly concerned about when imaging mitotic cells (Douthwright and Sluder, 2017; Laissue *et al*., 2017; Tosheva *et al*., 2020; Harada *et al*., 2023). Nevertheless, we did not repetitively image the same cell through multiple stages of mitosis to ensure that our study would not be influenced by even low levels of photo damage. To achieve the highest temporal resolution, and in light of the fact that both LLC-PK1 and PtK1 cells are known to remain relatively flat during mitosis(Rusan *et al*., 2001; Tirnauer *et al*., 2002; Wadsworth *et al*., 2005), we imaged a single z-plane of the cell over a 30-s duration using SDC.

LLC-PK1 cells were imaged on a Nikon Eclipse Ti microscope with a Nikon Plan Apo 100x/1.45-NA oil immersion objective. Images were captured with an ORCA-Fusion sCMOS camera using 488- and 561-nm solid-state lasers and standard filter sets.

PtK1 cells were imaged on a Nikon Ti microscope with a Nikon SR HP Apo TIRF 100x/1.49-NA oil immersion objective. Images were captured with a Photometrics Prime 95B sCMOS camera using 488- and 561-nm solid-state lasers and standard filter sets. PtK1 imaging was performed through use of the Vanderbilt Cell Imaging Shared Resource Nikon Center of Excellence.

### EB1 comet length analysis

EB1 comet lengths were quantified using a set of custom MATLAB (vR2022a; MathWorks) scripts as reported previously (Farmer*, Arpağ* *et al*., 2021; Cassidy*, Farmer* *et al*., 2024). Growth events were defined as periods of sustained growth lasting a minimum of 4 seconds (26 continuous frames), and growth rates were measured via linear regression of displacement over time. EB1-GFP intensity profiles were fitted to an exponential decay function, with the resulting decay constant used as a proxy measurement of EB1 comet length.

In brief, the start and end points of individual microtubule growth events were manually identified on kymographs, after which events shorter than 4 s were excluded by automated filtering. The initial estimate of microtubule tip position was estimated by assuming a constant growth rate between the first and last detected positions. Within each time frame, the brightest pixel corresponding to EB1 fluorescence was selected as the tip position within a ±5-pixel window centered on the initial estimate. The resulting series of tip positions was then fitted with a linear regression to calculate the growth rate of each event. To produce time-averaged intensity profiles, the tip positions identified in each frame of a given growth event were first aligned relative to the microtubule end (maximum intensity pixel), and the corresponding EB1 intensities were averaged across time to produce a single profile for each growth event. The microtubule lattice intensity was determined by averaging the fluorescence of five pixels along the lattice and then subtracting this value from the intensity of every pixel in the averaged profile for that segment.

EB1 comet length was determined by fitting the background-corrected intensity profile to an exponential decay function over 25 pixels, beginning with the pixel immediately following the assigned tip position. (Bieling *et al*., 2007; Farmer*, Arpağ* *et al*., 2021): I = Ae(−x/λ) + *B*, where *A* represents the intensity at the first pixel of the fit, *λ* corresponds to the comet decay length, and *B* corresponds to the y-offset. Omission of the 0th pixel intensity from the fit prevented any undetected subpixel variations in the microtubule tip structure from influencing the measured comet length. Growth events were then filtered to include only growth events with well-defined growth rates and where comet length could be confidently determined using R2 criterion of 0.8 and 0.9, respectively. Growth events that satisfied both R-squared criteria were grouped into 20 nm/s growth rate bins. To enable direct comparison of growth events across datasets, only data points within the overlapping range of growth rates (100-320 nm/s) were retained; data points falling outside this shared range were excluded from analysis. The binned data were then evaluated for outliers using an iterative Grubbs’ test, applied to each growth rate bin containing at least seven data points; bins containing less than seven data points, but at least three, were alternatively subjected to Grubbs outlier analysis. Following removal of outliers, any bins containing less than three growth events were removed. This dataset was then subject to relative error analysis. Briefly, for each growth event the relative error of the comet length was calculated (Relative error = Comet length CI/Comet length). The average relative error for each cell stage was then calculated. The cell cycle stage with the lowest average relative error (Interphase), was then set as the error threshold. Comets with a relative error greater than 2x the average relative error threshold were excluded from further analysis because these comet lengths could not be confidently determined. Weighted averages and weighted error for each bin were then calculated using 1/CI2 weighting. Averaged data points for each cell-cycle stage were plotted in GraphPad Prism and fit using unweighted least-squares linear regression. An extra sum-of-squares F-test was performed to determine whether the data from all stages could be adequately described by a single linear model. When the F-test indicated a significant difference (p < 0.05), the data were fit separately for each stage, with the y-intercept constrained to the best-fit y-intercept from the single-model fit, thereby enforcing a shared intercept across stages.

### Determination of cell cycle stage

To ensure the most controlled approach, we established a checklist of H2B-mCherry-18 and EB1-GFP fluorescence localization visual indicators that could be used to identify a given cell cycle stage (see examples in Figure 1A-F top images) (Kanda *et al*., 1998), in agreement with stage distinctions used by other studies (Sorce *et al*., 2015). Key indicators for each stage were as follows:

Interphase — Non-condensed chromatin with diffuse nuclear H2B-mCherry signal and an intact nuclear envelope. EB1-GFP signal was excluded from the nuclear region, and microtubules formed a radial array.

Prophase — Condensed chromosomes with an intact nuclear envelope. EB1-GFP signal remained excluded from the nuclear region.

Prometaphase — Condensed chromosomes and loss of the nuclear envelope, accompanied by diffuse EB1-GFP signal throughout the cell. Spindle assembly had initiated, as indicated by EB1 localization, but chromosomes were not yet aligned at the metaphase plate.

Metaphase — Chromosomes were aligned at the metaphase plate, and a bipolar spindle was fully formed.

Anaphase — Sister chromatids had separated and were no longer aligned at the metaphase plate, moving toward the spindle poles, which were visualized by EB1-GFP signal.

Two daughter cells — Two physically separated daughter cells were present and independent of one another. In some cases, a single-timepoint Z-stack was collected to confirm complete cytokinesis, demonstrating that EB1-GFP comets were spatially confined to their respective cells. Chromosomes had begun to, or were fully, decondensed.

## SUPPLEMENTARY FIGURES

**Figure S1.**
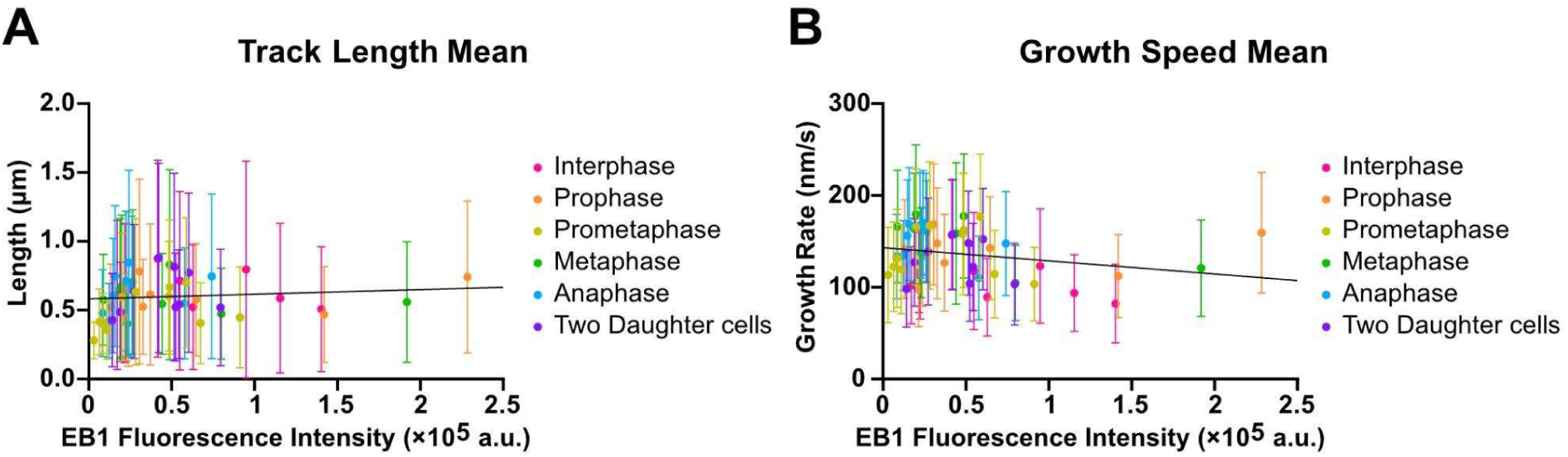
Microtubule growth parameters are not affected by EB1-GFP expression levels in LLC-PK1 cells. (A) Mean microtubule growth length plotted against corrected total cell fluorescence (CTCF) of EB1-GFP. (B) Mean microtubule growth speed plotted against CTCF of EB1-GFP. Microtubule dynamic parameters were quantified using plusTipTracker (Applegate et al., 2011). Each data point represents a single cell; error bars indicate standard deviation (SD). N = 56 cells. Linear regression analysis revealed no significant correlation between EB1-GFP expression level and either growth length (A, p = 0.26) or growth speed (B, p = 0.99).

**Video 1.** Video montage of EB1-GFP (magenta) and H2B-mCherry (cyan) in six different LLC-PK1 cells. Each of the investigated cell cycle stages are represented: top (left-to-right) interphase, prophase, prometaphase and bottom metaphase, anaphase, and two daughter cells.

## Abbreviations used

MAP: microtubule associated protein
SDC: spinning disc confocal
SD: standard deviation
CI: confidence interval

## Notes

### Competing Interest Statement

The authors have declared no competing interest.

